# Coupling environmental biosensors to inverter circuits in soil bacteria

**DOI:** 10.64898/2026.11.10.765548

**Authors:** Crislaine K. S. Rocha, Juan Rico, Ángel Goñi-Moreno

## Abstract

Genetic logic gates, such as inverting NOT functions, are routinely designed and optimised for a narrow set of laboratory-standard chemical inputs. This reliance limits their utility in real-world applications, where engineered chassis must navigate complex, niche-specific signals. Focusing on environmental synthetic biology, we engineered four biosensors responsive to ecologically relevant physical and chemical inputs (iron, oxygen, temperature, and light) and systematically coupled them to three repressor-based NOT logic gates (QacR, SrpR, and LitR) to generate 12 composite sensor–gate circuits. To evaluate cross-host robustness, all constructs were benchmarked in two non-model soil bacteria: *Pseudomonas putida* KT2440 and *Pseudomonas protegens* Pf-5. Our findings demonstrate that overall circuit performance is non-additive and cannot be predicted simply by summing individual sensor and gate dynamics. While standalone biosensors reliably detected their cognate inputs, concatenation across regulatory junctions introduced non-linearities, emergent dynamics, and host-dependent effects that caused several pairings to deviate unexpectedly from their predicted behaviour. Furthermore, host context directly governed logic execution: *P. protegens* Pf-5 exhibited superior overall circuit-processing capacity, whereas specific configurations, such as iron-responsive gates, performed better in *P. putida* KT2440. All 12 constructs are modularly integrated using the Standard European Vector Architecture (SEVA) format, providing a benchmarked toolkit to program complex information processing in real-world environments.

## Introduction

A central effort in synthetic biology^1^ since its beginning has been the engineering of computing functions.^2–5^ Foundational work predominantly focused on Boolean logic gates and circuits,^6,7^ but has also expanded to include analogue processing,^8,9^ neuromorphic networks,^10,11^ multicellular systems, ^12,13^ probabilistic computing,^14^ feedback controllers,^15^ and reservoir approaches, ^16^ among others. Using biological substrate for computation offers a compelling non-conventional computing route, using genetic networks to execute algorithmic structures that sense inputs and react to them in predefined ways to deliver outputs. While this input–algorithm–output pipeline reflects natural microbial behaviour, synthetic biology tools and methods enable the rational design of new-to-nature processes. Despite impressive progress across the field, using a small set of standard chemical inputs limits the application scope of such circuits—a problem that directly underpins the present study. Standard laboratory inducers, such as isopropyl *β*-D-1-thiogalactopyranoside (IPTG), anhydrotetracycline (aTc), or L-arabinose, serve as useful reference inputs for prototyping, inter-laboratory comparison and reproducibility. However, relying on these chemicals hinders real-world deployment, where living cells must operate within environments governed by inputs that differ vastly in their sensing mechanisms and dynamic performance.

Here, we focus on environmental applications^17,18^ as our target problem domain. The use of synthetic biology to address ecological challenges has been proposed extensively, ranging from advanced bioremediation strategies^19^ to large-scale ecosystem interventions. ^20,21^ Although significant technical and regulatory barriers remain, ^22,23^ applying cellular computing to tasks where natural microbes already excel as environmental engineers is both attractive and potentially transformative. ^24^ Characterising environmentally relevant inputs is therefore a key priority on this scientific agenda. Crucially, this problem domain demands not only specific input signals, but also appropriate cellular hosts. Parallel to the reliance on standard chemical inducers is the widespread use of the model organism *Escherichia coli*, despite growing evidence that host context profoundly impacts circuit performance. To address this, we employ two robust soil chassis: *Pseudomonas putida* KT2440^25,26^ and *Pseudomonas protegens* Pf-5.^27^ Together, domain-specific input signals and host chassis constitute the essential biological foundation for engineering genetic circuits tailored to real-world environments.

A substantial body of work has already focused on engineering biosensors:^28,29^ genetic devices capable of detecting target molecules or physical cues and triggering a corresponding cellular response. Numerous examples exist with direct relevance to ecological challenges, particularly in bioremediation, including biosensors for heavy metals,^30^ explosives,^31^ hydrocarbons,^32^ pesticides,^33^ and pharmaceuticals.^34^ However, biosensor development has historically progressed in parallel with genetic logic circuit design, with remarkably little crossover between the two fields. Coupling these distinct domains—interfacing real-world sensing devices directly with decision-making logic gates—is essential to reveal how genetic circuits actually perform under realistic dynamic conditions.

To address this challenge, we engineered four environmental biosensors responsive to iron, oxygen, temperature, and green light (Figure 1A), and coupled them to three NOT logic gates (inverters) based on the transcriptional repressors QacR, SrpR, and LitR, which were previously characterised in both target chassis (Figure 1B). By constructing a combinatorial library of all twelve sensor–NOT pairings in a modular format using the Standard European Vector Architecture (SEVA) and systematically evaluating their performance in both host strains (Figure 1C), we map the dynamic activity and operational limits of these environmental logic circuits. Crucially, the biological information encoded by the input signal was not always processed effectively across the downstream circuit elements, resulting in severe logic mismatches in specific pairings. Conversely, several sensor–gate combinations maintained robust logic performance, providing ideal, benchmarked candidates for future application in environmental synthetic biology.

**Figure 1:**
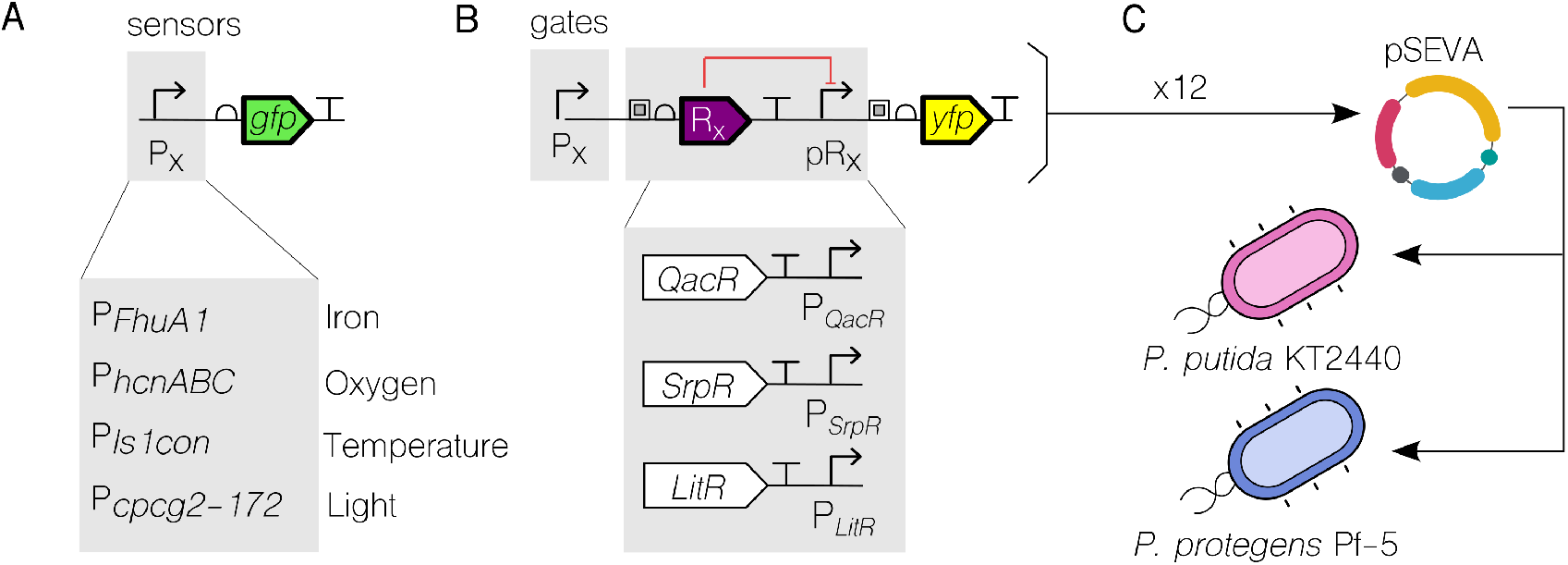
Pipeline for processing environmental inputs through modular biosensors. **A.** Biosensing promoters can be modularly exchanged within a standard SEVA vector architecture. These include promoters for sensing iron (*P_F huA_*_1_), oxygen (*P_hcnABC_*), temperature (*P_ls_*_1_*_con_*), and green light (*P_cpcg_*_2_*_−_*_172_). The reporter output for these standalone sensor constructs is green fluorescent protein (GFP). **B.** The genetic inverters (NOT logic gates) used in this study are based on the transcriptional repressors QacR, SrpR, and LitR. Sensor modules are positioned upstream of the repressor genes, driving the downstream reporter yellow fluorescent protein (YFP). **C.** Each of the resulting 12 sensor–NOT constructs was cloned into a SEVA plasmid backbone (featuring an RK2 origin of replication and a kanamycin resistance marker) and transformed into *Pseudomonas putida* KT2440 and *Pseudomonas protegens* Pf-5, yielding 24 engineered strains in total.

## Results

To facilitate direct performance comparisons among environmental sensing modules, all biosensors were standardised within an identical SEVA vector backbone, thereby minimising copy-number variability. Each sensing unit comprises an input-specific promoter placed upstream of a *gfp* reporter gene, enabling the quantitative monitoring of environmental responses. The cognate regulatory proteins that mediate signal detection were expressed either from the host’s native chromosomal loci or from independent synthetic constructs integrated into the chromosome. Decoupling these regulatory elements—keeping the sensor promoters on the plasmid while maintaining the signal detection machinery on the chromosome—maximises construct usability and versatility. Crucially, this modular design not only facilitates systematic characterisation, but also enables straightforward coupling to downstream genetic circuits.

### Chemical inputs for native NOT logic biosensing

The two chemical inputs evaluated in this study were iron and oxygen. Both biosensors exhibit intrinsic NOT logic: in the absence or at low levels of the input signal, reporter output is high, whereas high input concentrations leads to low output production. Despite sharing this inverse logic, the two sensors operate via distinct regulatory mechanisms and internal architectures.

In the case of iron (Figure 2A), the biosensor was developed based on the ferric uptake regulator, Fur. This regulatory system coordinates gene expression in response to bacterial iron availability.^35^ A set of iron-responsive, Fur-regulated promoters was previously characterised to identify optimal induction kinetics, establishing *P*_FhuA1_ as the most efficient candidate^36^—and thus the promoter chosen for this study. To preserve physiological regulation, Fur was not overexpressed, relying instead on its native chromosomal expression level at both hosts. In the absence of iron, Fur is unable to repress gene expression, resulting in high GFP output. Under iron-rich conditions, Fur binds divalent iron (Fe^2+^), forms an active complex, and interacts with the Fur-box operator present within the *P*_FhuA1_ promoter, thereby repressing transcription.

**Figure 2:**
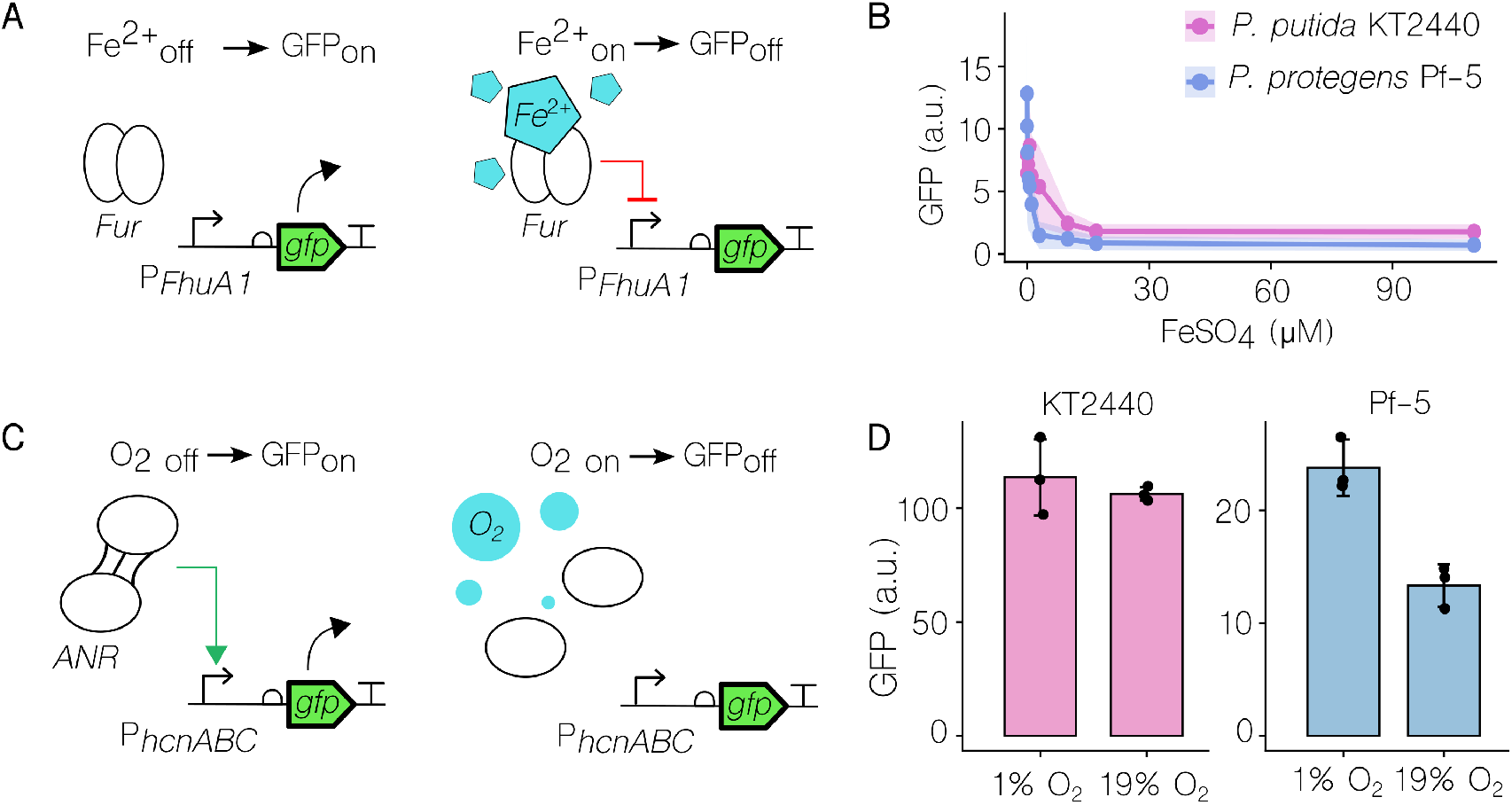
Characterisation of biosensors with native NOT logic for chemical signals: iron and oxygen. **A.** Mechanism of the iron biosensor based on the *P_F huA_*_1_ promoter. This promoter is constitutively active by default, driving GFP expression in the absence of Fe^2+^. When Fe^2+^ is present in the growth medium, it binds to the endogenously expressed ferric uptake regulator (Fur) to form a complex that represses promoter activity. **B.** Iron biosensor transfer functions measured by GFP fluorescence in response to increasing FeSO_4_ concentrations in *Pseudomonas putida* KT2440 and *Pseudomonas protegens* Pf-5. The response follows an intrinsic NOT logic, where increasing input signal leads to a decrease in reporter output. **C.** Mechanism of the oxygen biosensor based on the *P_hcnABC_*promoter. Under low oxygen availability, the transcriptional regulator ANR forms an active complex that induces promoter activity and GFP expression. Exposure to high oxygen concentrations leads to the disassembly of the active ANR complex, stopping promoter induction. **D.** Oxygen biosensor performance across low (1%) and high (19%) oxygen conditions in both host strains, displaying a subtle NOT-like response with lower reporter output under high input concentrations.

To characterize iron sensing, reporter activity was measured in iron-free minimal medium (M9-glucose) supplemented with increasing concentrations of FeSO_4_ across both bacterial hosts (Figure 2B). Although the biosensor responded to iron availability via NOT logic in both strains as expected, the transfer function exhibited differing dynamic characteristics between hosts. *P. protegens* Pf-5 demonstrated a higher baseline output in the absence of iron and greater sensitivity to low signal concentrations, with reporter expression dropping sharply at low iron concentrations and resulting in a more digital, switch-like response curve. In contrast, *Pseudomonas putida* KT2440 displayed a distinct response profile marked by lower maximum output expression and a more gradual transition towards baseline. These discrepancies likely reflect host-dependent variations in iron uptake kinetics and Fur-mediated regulation, underscoring the influence of chassis context on biosensor performance. Notably, soluble iron concentrations in soil solutions typically range from a few micromolar up to 200 *µ*M,^37^ confirming that the sensitivity of this sensor is well matched to environmental applications. To further assess the response of the sensor to iron availability, *P. protegens* Pf-5 carrying *P*_FhuA1_ report was exposed to increasing amount of the iron chelator 2,2’-dipyridyl (DPD). DPD induction increased GFP expression, consistent with relief of Fur-mediated repression under decreasing iron availability (Supplementary Figure S1).

In the case of oxygen (Figure 2C), we selected the *P*_hcnABC_ promoter, whose response under oxygen-limiting conditions is regulated by ANR, an FNR-family transcriptional factor that responds to oxygen availability in *Pseudomonas*.^38,39^ *P*_hcnABC_ was chosen from a set of candidate oxygen-sensing promoters based on its performance in a initial screening (Supplementary Figure S2). As with the iron biosensor, this oxygen-sensing system relies on the host’s native regulatory machinery rather than heterologous overexpression. Under low O_2_conditions, active ANR efficiently induces *P*_hcnABC_, driving high reporter gene expression. Conversely, under high O_2_ conditions, molecular oxygen disrupts the active [4Fe *−* 4S]^2+^ iron–sulfur cluster of the ANR complex, abolishing its DNA-binding capacity and returning reporter expression to baseline levels.

The biosensor performance (Figure 2D) was tested with 1% oxygen as the low input state and 19% oxygen as the high input state, resulting in very different behaviour in both chassis. In KT2440 maximum output expression levels were 5-fold higher than in Pf-5. But only in Pf-5 the separation between output states (i.e., different output low vs high states) was statistically significant (see Methods). Although this state separation in KT2440 timidly shows a trend towards proper NOT function, there is wide variability and noise across replicates to send a definitive message. It is therefore in Pf-5 where the biosensor performs best. This suggests that finding the appropriate chassis for a given function may become crucial when moving away from pristine laboratory proofs of concept towards niche-specific applications.

### Physical inputs for native YES logic biosensing

The two physical input signals included in this study were temperature and light. While their mechanistic specifications are radically different, they both share a YES logic function: at low input signal, the output should be also low; at high input signal, high output expression is expected.

Regarding the temperature biosensor (Figure 3A), we selected the *P*_ls1con_ promoter, which is repressed by the temperature-sensitive *λ* bacteriophage repressor cI857. The cassette expressing cI857 was synthesised and integrated into the chromosome of both *Pseudomonas* strains, yielding engineered hosts capable of responding dynamically to temperature shifts. At low temperatures, cI857 actively binds the promoter operator sites to repress reporter expression, establishing a low-output state in response to a low-input signal. Conversely, at elevated temperatures, cI857 loses its DNA-binding capability, relieving transcriptional repression and allowing *P*_ls1con_ to drive reporter expression.

**Figure 3:**
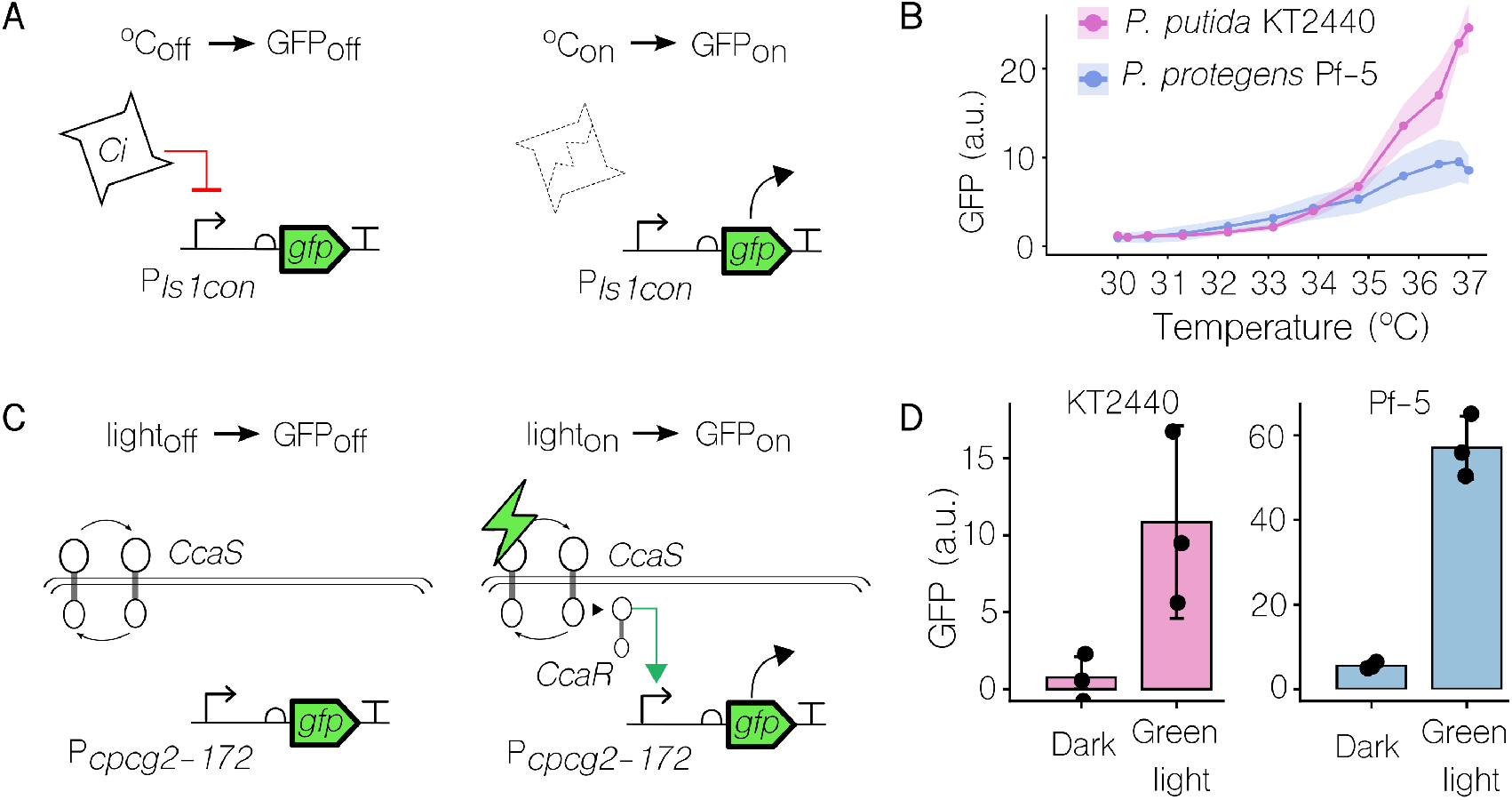
Characterisation of biosensors with native YES logic for physical signals: temperature and light. **A.** Regulatory mechanism of the temperature-sensing module based on the promoter *P*_ls1con_, controlled by the temperature-sensitive *λ* bacteriophage

To characterise the performance of the temperature biosensor, cultures were incubated across a thermal gradient ranging from 30 *^◦^*C to 37 *^◦^*C (Figure 3B). Both strains displayed a clear temperature-dependent increase in GFP expression across the evaluated range, whereas repressor cI857. At low temperatures (30 *^◦^*C), cI857 represses transcription. At elevated temperatures (*≥* 37 *^◦^*C), cI857 loses DNA-binding activity, resulting in derepression of *gfp* expression. **B.** Temperature-dependent response curves in *Pseudomonas putida* KT2440 and *Pseudomonas protegens* Pf-5 across a temperature gradient (30 *^◦^*C to 37 *^◦^*C), demonstrating gradual activation following native YES logic. **C.** Schematic of the green-light sensing module based on the CcaS/CcaR two-component optogenetic system. In the dark (light_off_), reporter expression remains at basal levels. Upon green-light illumination (520 nm), the sensor kinase CcaS autophosphorylates and transfers phosphate to the response regulator CcaR, which activates transcription from promoter *P*_cpcg2-172_. The chromophore phycocyanobilin required for photoreceptor function is synthesized via co-expressed Ho1 and PcyA enzymes. **D.** Reporter output in *P. putida* KT2440 and *P. protegens* Pf-5 under dark and green-light conditions, showing robust light-dependent induction in both hosts cells. control strains lacking the cI857 repressor exhibited temperature-independent expression profiles (Supplementary Figure S3), confirming that regulation was specifically mediated by the engineered thermosensing mechanism. While *Pseudomonas putida* KT2440 and *Pseudomonas protegens* Pf-5 exhibited virtually identical induction profiles up to 35 *^◦^*C, their performance diverged significantly at higher temperatures, with *P. putida* KT2440 reach ing substantially higher reporter output levels at 37 *^◦^*C. Although this strong expression might initially suggest that *P. putida* KT2440 is the superior chassis for temperature sensing, downstream coupling of this biosensor to genetic NOT gates yielded markedly better circuit performance in *P. protegens* Pf-5—a striking and counter-intuitive phenomenon that is discussed further in Figure 4.

**Figure 4:**
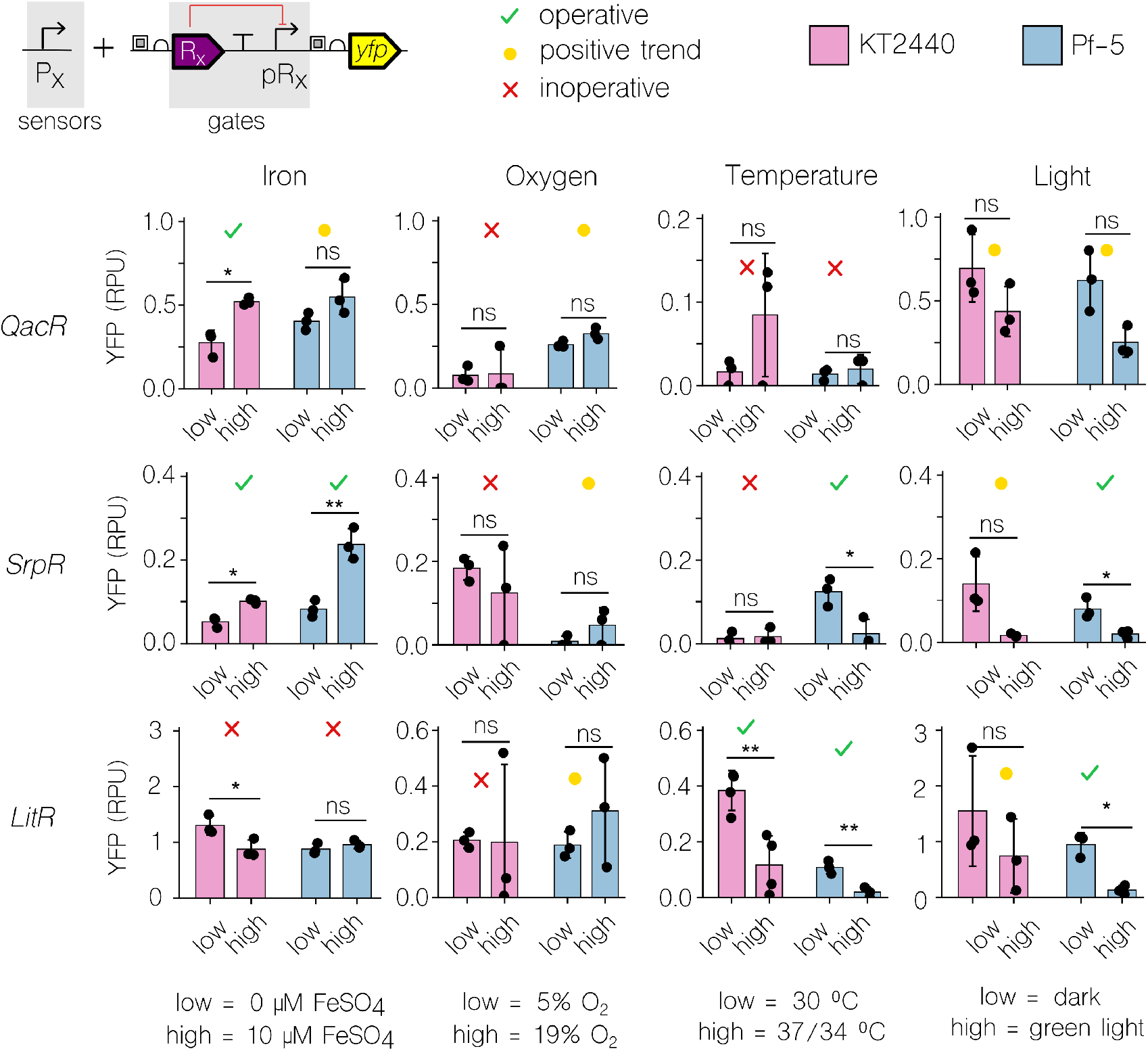
Functional performance of environmental biosensors coupled to genetic NOT logic gates. Each of the four environmental biosensors (iron, oxygen, temperature, and green light) was concatenated upstream of three distinct repressor-based NOT gates (QacR, SrpR, and LitR). Bar plots compare lowversus high-input signal responses across *Pseudomonas putida* KT2440 (pink) and *Pseudomonas protegens* Pf-5 (blue), measuring yellow fluorescent protein output (YFP, arbitrary units). Input ranges were defined as follows: iron (0 vs. 10 *µ*M FeSO_4_), oxygen (5% vs. 19% O_2_), temperature (30 *^◦^*C vs. 37 *^◦^*C for KT2440 and 34 *^◦^*C for Pf-5), and green light (dark vs. green light illumination). Overhead symbols classify circuit operational status: green tick = fully operative function; yellow circle = positive qualitative trend without statistical significance; red cross = inoperative/incorrect behaviour. Statistical significance between low and high states was quantified as *ns*: nonsignificant (p *≥* 0.05); * (p < 0,05): significant; *∗∗* (p < 0,01): very significant.

Regarding the green light biosensor (Figure 3C), we integrated the heterologous CcaS/CcaR optogenetic system into the chromosome of both host strains. This architecture was adapted from a previously reported construct in which both the core two-component machinery and the cognate promoter resided on a plasmid; in our implementation, only the target *P*_cpcg2-172_ promoter is carried on the SEVA plasmid backbone. ^40^ In the dark, the promoter remains inactive and reporter output is only expressed at low basal levels. Upon illumination with green light, the optogenetic sensor phosphorylates the response regulator to induce transcription from *P*_cpcg2-172_, driving robust reporter expression. The system thus executes intrinsic YES logic, converting low-input to low-output and high-input to high-output states.

The performance of the green light biosensor (Figure 3D) exhibited similar qualitative trends in both host chassis, characterised by a clear separation between dark (low) and illuminated (high) states upon exposure to a 520 nm green light source, alongside notable inter-replicate variability in the high (on) state. Green-light induction reached 14-fold over the dark state in *P. putida* KT2440 and 10-fold in *P. protegens* Pf-5. Despite this slightly lower fold induction, Pf-5 achieved substantially higher maximum reporter expression levels than KT2440 (approximately 60 versus 10 arbitrary units, respectively), whereas KT2440 exhibited lower basal expression in the dark. Because reporter constructs, instrument settings, and flow cytometry acquisition protocols were kept identical across experiments, these dynamic output levels are directly comparable. In this case, this higher expression levels at high input signal impacts positively on downstream circuit performance when coupled to logic gates, as evaluated below.

### Performance of sensor–gate pairings

The coupling of environmental biosensors to genetic NOT gates enabled the systematic study of information processing within composite genetic circuits. Sensor promoters were positioned upstream of repressor-based gates in a combinatorial layout: each of the four biosensors was paired with three distinct NOT logic gates (QacR, SrpR, and LitR) to generate 12 unique sensor–gate constructs. The circuits output performance was quantified via the downstream YFP reporter of the gate. The performance metrics across both host strains are presented in Figure 4.

To establish functional performance metrics, discrete low- and high-input signal values were chosen based on the dynamic range observed for each standalone biosensor (Figures 2 and 3). These input regimes were defined as: 0 vs. 10 *µ*M FeSO_4_ for iron, 5% vs. 19% O_2_ for oxygen, 30 *^◦^*C vs. 37 *^◦^*C (for *Pseudomonas putida* KT2440) and 34 *^◦^*C (for *Pseudomonas protegens* Pf-5) for temperature, and dark vs. green-light illumination for the optogenetic system. The output states resulting from these high and low inputs were evaluated to assess the functional logic of each sensor–gate pairing.

Although the architecture of both the individual biosensors and the logic gates is relatively straightforward, and their standalone performance would suggest smooth logic execu- tion across all pairings, empirical evaluation revealed a more complex reality. While several constructs exhibited fully operative logic, specific sensor–gate combinations displayed unexpected or counter-intuitive behaviour.

Since the iron biosensor functions natively as a NOT gate, its concatenation with a downstream inverter logic gate was expected to invert the response, yielding an overall YES logic function. For the SrpR gate, this predicted YES logic was achieved optimally in both hosts, displaying a statistically significant output induction, albeit with lower overall expression levels in *P. putida* KT2440 compared to *P. protegens* Pf-5. When the inverter module was replaced with QacR, statistically significant YES logic switching was observed exclusively in *P. putida* KT2440, although *P. protegens* Pf-5 exhibited a clear qualitative trend towards correct circuit operation. Conversely, the LitR gate was completely unable to process the iron biosensor inputs, rendering the resulting sensor–gate pair inoperative in both host strains. Given that the LitR gate functions reliably when driven by an IPTG-inducible promoter (Supplementary Figure S4), this failure points to emergent, system-level dynamics or promoter–repressor context dependencies that would require future further investigation. The second column in Figure 4 summarizes the oxygen–coupled circuits, which likewise were expected to execute overall YES logic. These constructs exhibited the lowest success rate across the entire dataset—an outcome that aligns with the performance of the standalone oxygen biosensor, which demonstrated narrow separation between low- and high-input states, particularly in *P. putida* KT2440. In this host chassis, none of the three oxygen–gate combinations functioned as expected. Conversely, in *P. protegens*Pf-5, all three circuits displayed a consistent qualitative trend towards expected logic implementation, although these differences did not achieve statistical significance under our evaluated metrics (see Methods). This observation indicates that downstream signal processing can, to a degree, buffer and propagate marginal differences in initial sensing. Furthermore, in *P. protegens* Pf-5, all three repressor gates performed comparably—including LitR, which did not exhibit the disruptive context-dependent constraints observed in the iron–LitR pairing.

The third and fourth columns in Figure 4 represent circuits coupled to natively YES-logic sensors (temperature and green light, respectively), which were expected to execute overall NOT logic upon concatenation with a downstream inverter gate. Regarding the temperature biosensor, it is particularly noteworthy that, although the standalone sensor exhibited superior dynamic range and separation in *P. putida* KT2440 (Figure 3B), the complete sensor–gate circuits achieved significantly higher performance in *P. protegens* Pf-5—most notably when paired with the SrpR and LitR gates. This striking divergence underscores that circuit viability depends not merely on initial input sensing, but on how signal dynamics are processed across genetic nodes; counter-intuitively, a larger input dynamic range does not inherently guarantee optimal downstream gate performance. Furthermore, underlying contextual constraints proved limiting for the QacR gate, where both host chassis failed to deliver functional NOT logic switching.

Lastly, the green-light biosensor yielded high operational success across all tested NOT gates and host strains. Performance was broadly comparable between the two chassis—a surprising result given that the standalone optogenetic sensor generated substantially higher reporter expression in *P. protegens* Pf-5 (Figure 3D). Evidently, these disparities in input magnitude are attenuated as the signal is processed downstream towards the ultimate YFP output. Indeed, all light–gate pairings displayed qualitative operational behaviour, though statistically significant switching was achieved exclusively in *P. protegens* Pf-5. While this enhanced statistical confidence may stem from the larger initial separation between low- and high-input signal states in this host, this hypothesis remains unclear.

## Discussion

This study represents a systematic effort to tailor genetic logic circuitry to real-world niche applications in environmental synthetic biology. By replacing standard laboratory chemical inputs with ecologically relevant environmental signals (iron, oxygen, temperature, and light) across a combinatorial set of repressor-based NOT logic gates (QacR, SrpR, and LitR), we evaluated 12 composite sensor–gate pairings. Crucially, by extending this pipeline to two environmentally relevant soil bacteria—*Pseudomonas putida* KT2440 and *Pseudomonas protegens* Pf-5—we provide a comprehensive characterisation of chassis-dependent circuit performance in non-model hosts.

A major conclusion of this work is that host-dependent contextual features ^41^ serve as a primary determinant of circuit viability, in line with previous observations on the context dependencies of genetic inverters.^7,42,43^ Functional behaviour was highly host-specific: while iron-sensing circuits exhibited superior performance in *P. putida* KT2440, green-light processing achieved higher expression and statistical confidence in *P. protegens* Pf-5. To our knowledge, this work also provides the first CcaS/CcaR optogenetic sensor in *P. protegens* Pf-5, extending light-control expression, so far shown in *P. putida* KT2440,^40^ to a second soil *Pseudomonas* chassis. Overall, *P. protegens* Pf-5 demonstrated superior circuit-processing capacity across the 12-construct matrix, yielding higher success rates and operational reliability. Such disparities are unsurprising given that soil microbes evolved distinct metabolic, regulatory, and uptake machinery to adapt to specific micro-environmental niches (e.g., differential Fur-mediated iron regulation^35^ or oxygen-responsive ANR cascades^38,39^). Consequently, exploiting non-model chassis and quantifying host–circuit interactions is not merely beneficial, but essential for application-driven synthetic biology.

Furthermore, our quantitative dataset explicitly demonstrates that signal sensing and downstream logic processing are decoupled phenomena. Standalone biosensor efficiency showed no direct correlation with successful downstream logic execution. Instead, sensorgate concatenation induced non-linear, emergent dynamics heavily dependent on cellular ^44^ and genetic context.^45^ A prime example is the temperature biosensor: although it displayed a far superior dynamic range in *P. putida* KT2440 as an isolated module, its concatenation with inverter gates failed to yield functional logic in this host, whereas *P. protegens* Pf-5 successfully executed the expected NOT logic. Similarly, promoter–repressor context dependencies led to specific gate failures, such as the complete operational breakdown of LitR when coupled to iron sensing in both strains, despite LitR functioning reliably under IPTG control.^46,47^ Conversely, in marginal sensors such as oxygen, signal propagation through downstream gates partially buffered weak input separation in *P. protegens* Pf-5.

These idiosyncratic, pair-specific behaviours indicate that universal design rules cannot yet be reliably inferred for composite environmental circuits. Nevertheless, this expands the repertoire of functional behaviours accessible across diverse chassis. All constructs in this study were standardised using the Standard European Vector Architecture (SEVA) format,^48^ establishing a robust, modular toolkit ready to be used in environmental synthetic biology and bioremediation projects.

## Materials and Methods

### Strains, media, growth conditions

In this work *Escherichia coli* strains were used for cloning and plasmid propagation, whereas *P. putida* KT2440 and *P. protegens* Pf-5 were used for biosensor and genetic circuit characterisations (Table S1). Unless otherwise stated, bacterial cultures were grown in lysogeny broth (LB) supplemented with appropriate antibiotics (50 *µ*g/mL of kanamycin, 10 *µ*g/mL of gentamicin, 150 *µ*g/mL of ampicillin, and/or 25 *µ*g/mL of chloramphenicol). M9 minimal medium supplemented with 0.2% citrate was used for conjugation procedures, whereas M9 supplemented with glucose (20 mM) was used for iron biosensor and iron-responsive circuit characterisation. Liquid cultures were routinely incubated with shaking at 200 rpm. *E. coli* cultures were grown at 37°C and Pseudomonas at 30°C, unless otherwise specified for individual assays.

### Cloning design

Genetic constructs were assembled following the Golden Standard modular cloning framework (^49^). DNA parts were obtained by synthesis, PCR amplifications with Phusion High-Fidelity DNA polymerase (Thermo Fisher Scientific, Waltham, Massachusetts), or from the Golden Standard part library. PCR primers were designed to introduce *Bsa*I-compatible sequences for modularization and, when required, *Eco*RI/*Hin*dIII restriction sites for conventional cloning into pSEVA182 (Table S3). For conventional cloning, vectors were doubledigested and dephosphorylated using enzymes from New England Biolabs (NEB, Ipswich, MA, USA). Level 1 constructs were assembled using pSEVA2219-g1 or pSEVA2219-tn7-g1 as recipient vectors. Modular cloning reactions contained 50 ng of vector, 50 ng of each required DNA part, 20 U of *Bsa*I (NEB), 20 U of T4 DNA ligase (Promega, Madison, WI, USA), T4 ligase buffer, and 0.1 mg/mL bovine serum albumin (BSA). Reactions were incubated for 25 cycles of 4 min at 37 °C and 4 min at 16 °C, followed by 5 min at 37 °C and 20 min at 80 °C. Five microliters of the assembly reaction were used to transform chemically competent *E. coli* TOP10 or DH5*α* cells as described by Inoue et al. (^50^). Plasmid DNA was purified using the Monarch Plasmid Miniprep Kit (NEB), and PCR products were purified using the Monarch PCR & DNA Cleanup Kit (NEB). Oligonucleotides were obtained from Integrated DNA Technologies (IDT, Coralville, IA, USA). All final constructs were verified by diagnostic PCR and DNA sequencing (Plasmidsaurus, Eugene, OR, USA).

### Construction of biosensors, NOT gates, and reference constructs

Environmental biosensors were assembled using a common modular architecture comprising the corresponding input-responsive promoter, BCD12 ribosome-binding site, domesticated fluorescent GFP reporter, and rpoCD transcriptional terminator in a pSEVA2219-derived backbone. The sensing promoter was exchanged according to the environmental input under study, while the remaining architecture was maintained whenever possible. NOT-gate circuits were generated by adapting previously developed LacI/IPTG-responsive inverter architectures.^46,47^ The original Ptac/LacI input module was removed by PCR amplification of the remaining inverter cassette using primers introducing Golden Standard B-G assembly overhangs. This strategy preserved the downstream inverter architecture while allowing different environmental sensing promoters, assembled as A–B modules, to be introduced at the input. The resulting circuits therefore contained an environmental sensing promoter controlling the corresponding QacR-, SrpR-, or LitR-based inverter, followed by the cognate repressor-regulated output promoter and YFP reporter. To enable direct comparison between biosensor and NOT-gate outputs, corresponding biosensor constructs were also generated using YFP as reporter but lacking the inverter module. A constitutive YFP reference construct was assembled using the J23101 promoter together with the corresponding insulator, RBS, YFP reporter, and terminator in the same pSEVA-derived architecture and was used for calculation of relative promoter units (RPU). In experiments requiring an additional constitutive positive fluorescence control, a GFP-expressing construct driven by the Bba_J23106 promoter was used.

### Chromosomal integration of regulatory modules

Regulatory modules required for temperature- and light-responsive sensing were integrated into the chromosome using a Tn7-based system. For temperature sensing, the CIts regulatory cassette was PCR-amplified from a pre-existing construct from the laboratory collection and assembled into a Level 1 vector containing the elements required for Tn7-mediated chromosomal integration. The CIts module was placed under the control of a constitutive Anderson promoter (Bba_J23106) and BCD12 RBS, followed by the rpoCD terminator. For green-light sensing, the regulatory module was derived from pGreenL. ^40^ To minimize repeated DNA sequences within the final cassette, the promoter and terminator associated with *ccaS* were replaced by Bba_J23106 and the *rnpB* terminator, respectively, whereas the terminator associated with *ccaR* was replaced by rpoCD. The individual components of the light-responsive regulatory system, including *ccaS*, *ccaR*, and the chromophore-biosynthesis module, were assembled by Golden Standard MoClo and subsequently cloned into a Tn7-compatible Level 1 vector. The resulting Tn7 regulatory constructs were introduced into *P. putida* KT2440 and *P. protegens* Pf-5 using a Tn7-mediated conjugation protocol as previously described.^51,52^ Correct insertion at the chromosomal attTn7 site adjacent to *glmS* was verified by diagnostic PCR and sequencing of the corresponding genomic locus. Additional diagnostic PCR assays using the PS3-PS6 primer sets were performed to screen for potential ectopic integration events. Strains carrying the verified chromosomal integrations were subsequently used as recipients for transfer of the biosensor and NOT-gate plasmids by triparental mating (^47,52^).

### Characterisation of the biosensors

**Iron biosensor**: Overnight cultures were grown in M9-glucose medium supplemented with kanamycin. Saturated cultures were diluted in fresh M9-glucose-Km medium to an initial OD_600_ of 0.1 and incubated for 3 h at 30°C with shaking at 200 rpm, in order to achieve log phase. Subsequently, 180 *µ*L of culture were transferred to 96-well plates in technical duplicate, and 20 *µ*L of FeSO_4_ (iron(II) sulfate heptahydrate, Acros Organics, Geel, Belgium) solution was added to obtain a final concentration of 0, 0.01, 0.1, 0.3, 0.6, 1.0, 3, 10, 17, and 110 *µ*M in a final volume of 200 *µ*L. Plates were incubated for 22 h at 30°C with shaking at 200 rpm. At the end of the incubation period, OD_600_ and fluorescence were measured using Varioskan*^TM;^*LUX multimode microplate reader (Thermo Fisher Scientific, Waltham, Massachusetts), and samples were additionally analysed by flow cytometer. In parallel, iron limitation was induced in *P. protegens* Pf-5 grown in LB-medium by addition of the iron chelator 2,2’-Dipyridyl (DPD, Thermo Fisher Scientific, Waltham, Massachusetts). The same experimental procedure described above was followed, replacing FeSO_4_ supplementation with DPD at final concentrations of 0, 50, 100, 200, 300, 400, 600, 800, and 1000*µ*M.

### Oxygen biosensor

Overnight cultures were grown in LB medium supplemented with kanamycin. Saturated cultures were diluted directly into 96-well plates to an initial OD_600_ of 0.01 and a final volume of 200 µL per well, with two technical replicates per condition. A part of the auto-fluorescence control strain carrying the pSEVA2219 empty plasmid, a positive control containing GFP under the constitutive Bba_J23106 promoter was also included. Plates were incubated at 30°C in a Varioskan*^TM^*LUX plate reader equipped with the integrated gas-control module. Oxygen concentration was controlled by supplying nitrogen (N_2_) to the instrument with a range from 1 to 21%. Cultures were first incubated for 16 h under 18-19% *O*_2_, followed by an additional incubation for 4 h under 1% O_2_. Fluorescence and growth were monitored after every 20 min during all experiment. Results were based on endpoint measurements after 4 h of exposure of 1% *O*_2_.

### Temperature sensor

Overnight cultures were grown in LB medium supplemented with kanamycin and gentamicin. 100 *µ*L of saturated cultures were transferred to 96-well PCR plate. Prior to use, the plate was exposed to UV light for 15 minutes in a laminar airflow (LAF) bench. The plate was sealed with a sealing film for microplates (BRAND GmbH + Co KG, Wertheim, Germany), and incubated at the Mastercycler Nexus Gradient thermocycler (Eppendorf, Hamburg, Germany) for 10 hours, programmed with a temperature gradient: 30, 30.2, 30.6, 31.3, 32.2, 33.1, 33.9, 34.8, 35.7, 36.4, 36.8, 37 °C. After incubation, samples were diluted 1:1 with 0.22 µm filtered LB medium and transferred to a standard 96-well plate for flow cytometry analysis. In addition to the auto-fluorescence control strain, the wild type strains lacking the integrated temperature Ci regulator but carrying the biosensor plasmid were included as a regulatory control.

### Green-light sensor

Overnight cultures were grown in LB medium supplemented with kanamycin and gentamicin. Saturated cultures were diluted in fresh LB medium to an initial OD_600_ of 0.1 and distributed in technical duplicates into two identical 96-well plates. One plate was completely covered with foam to prevent light exposure and served as the dark condition, whereas the second plate was left uncovered and exposed to green light. Green light illumination (approximately 530 nm) was provided using a custom-built 3D-printed light box lined with a commercial 5050 RGB LED strip (90 LEDs, 3 m; OYA Smart Lighting (Shenzhen) Co., Ltd., Shenzhen, China). The light box was placed inside a Fisherbrand Incubating Mini Shaker (Thermo Fisher Scientific, Waltham, Massachusetts) for 22 h at 30°C. Samples OD_600_ and fluorescence were analysed at the end of the incubation using a Varioskan*^T M^* LUX plate reader. In addition to the auto-fluorescence control strain, the wild type strains lacking the integrated light-regulatory module but carrying the biosensor plasmid were included as a regulatory control.

### Characterisation of sensor–gate pairings

QacR-, SrpR-, and LitR-based logic gate circuits were evaluated in combination with the corresponding environmental sensing modules under the conditions established for each sensor, with assay-specific modifications described below. In each assay, the three sensor-gate pairing were analysed in parallel with the corresponding host carrying an empty pSEVA2219 vector as an auto-fluorescence control, a constitutive Bba_J23101-YFP reference construct used for RPU-normalization, and the corresponding environmental biosensor carrying YFP as a reporter.

### Iron-logic circuit

Iron-sensing logic gates were characterized using the same general workflow described for the biosensor, but using a reduced FeSO_4_ concentration range (0, 0.1, 1.0, 10, 110 *µ*M).

### Oxygen-logic circuit

In contrast with oxygen biosensor characterisation, the lowoxygen condition for sensor-gate pairing was set to 5% O_2_. This concentration was selected as a less restrictive low-oxygen condition, with the aim at supporting culture growth while maintaining the gate output in the expected low state. Overnight cultures grown in LB medium were diluted to an initial OD_600_ of 0.1 in 96-well plates and incubated for 10 h under either low-oxygen (5% O_2_) or high-oxygen conditions [18–19% O_2_]. Values reported for the sensor-gate pairing characterisation correspond to measurements obtained at the 10 h time point.

### Temperature-logic circuit

In contrast with the biosensor characterisation, an additional pre-culture step was included before incubation under temperature gradient. Following overnight growth, cultures were diluted in fresh LB medium at initial OD_600_ 0.1 and grown for 2 h at 30 °C with shaking at 200 rpm. Cultures were then transferred to 96-well PCR plates and subjected to the temperature gradient as described above.

### Green light-logic circuit

The same experimental setup described for the green-light biosensor was used for the sensor-gate pairings.

### Fluorescence measurements and data processing

Bulk fluorescence and OD_600_ were measured using the Varioskan*^TM^*LUX plate reader. Platereader measurements were blank-corrected prior to normalization. Fluorescence values were normalised to OD_600_, and host auto-fluorescence was corrected by subtracting the signal obtained from the corresponding empty-vector control. Flow cytometry analyses were performed using a Miltenyi Biotec MACSQuant VYB flow cytometer (Miltenyi Biotec), acquiring 10,000 singlet events per sample using the B1 channel. The instrument was calibrated using MACSQuant Calibration Beads, following the general acquisition procedure described previously (^47^). Data was processed by MACSquantify software (Version 2.13.3), and auto-fluorescence was corrected as described above. Biosensors responses were expressed as background-correlated GFP fluorescence in arbitrary units (a.u.). For sensor–gate pairing experiments, YFP fluorescence from the sensor–gate circuits and the corresponding YFP biosensor controls was normalised to the constitutive Bba_J23101-YFP reference and expressed as RPU. RPU values were calculated from the corresponding fluorescence measurements as:

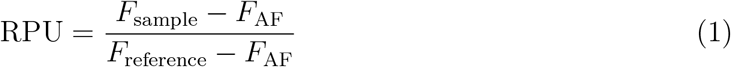

Where *F*_sample_ is the fluorescence of the analysed strain, *F*_AF_ is the fluorescence of the auto-fluorescence (empty vector) control strain, and *F*_reference_ is the fluorescence of the strain carrying the YFP controlled by Bba_J23101 constitutive promoter.

### Statistical analysis

Technical replicates were averaged before statistical analysis, such that each independent experiment contributed a single value per experimental condition. Unless otherwise indicated, experiments were performed using three independent biological replicates; exceptions are specified in the corresponding figure legends. Data are presented as individual biological replicates together with the mean and standard deviation (SD). Statistical comparisons between two experimental conditions were performed separately for each bacterial host. Welch’s unpaired two-sample *t* -test was used for comparisons between independent experimental conditions. For temperature experiments in which the same biological culture was distributed across the different temperature conditions, paired *t* -tests were used for endpoint comparisons. For the green-light regulatory-control experiment, a two-way ANOVA was used to evaluate the effects of regulatory construct, light condition, and their interaction. Statistical analyses were performed in R (version 4.5.2) using RStudio (version 2026.09.0, Build 174; Posit Software, PBC). A *p*-value < 0.05 was considered statistically significant. Significance levels were represented as ns (*p ≥* 0.05), * (*p* < 0.05), ** (*p* < 0.01), and *** (*p* < 0.001).

## Data availability

The dataset generated and analysed in this study are publicly available at: https://saco.csic.es/s/8YH7z5eRFSwdmTn. GenBank accession numbers for each construct generated in this study are provided in the supplementary table S2.

## Author contributions

CKSR: investigation, methodology, formal analysis, data curation, original draft; JR: methodology, formal analysis, data curation; AGM: conceptualization, supervision, writing. All authors contributed to the review and editing of the manuscript.

## Acknowledgements

This work was supported by the following funding sources: ECCO (ERC-2021-COG-101044360) and NEWTON (ERC-2025-POC-101248112) contracts of the EU, and grants MULTISYN-BIO (PID2023-152470NB-I00) and ADA LOVELACE IN VIVO (AIA2025-164235-C44) funded by MICIU/AEI/ 10.13039/501100011033.

## Supporting information

The Supporting Information includes a PDF file containing supplementary figures with additional experimental characterisation and supporting methodological details, as well as supplementary tables providing information on strains, plasmids, and primer sequences.

