## Supplementary for "Coupling environmental biosensors to inverter circuits in soil bacteria"

### Supplementary Information to: Coupling environmental biosensors to inverter circuits in soil bacteria

### Contents


- Additional characterisation of environmental biosensors and regulatory modules
- Supplementary Figure S1. Iron biosensor characterisation using DPD
- Supplementary Figure S2. Screening of oxygen-responsive promoters
- Supplementary Figure S3. Regulatory controls for the temperature- and green-light-responsive systems
- Supplementary Figure S4. Reference response of the QacR-, SrpR-, and LitR-based NOT gates to IPTG
- Supplementary Table S1. List of bacterial strains used in this study
- Supplementary Table S2. List of plasmids and genetic constructs used in this study
- Supplementary Table S3. List of primers used in this study

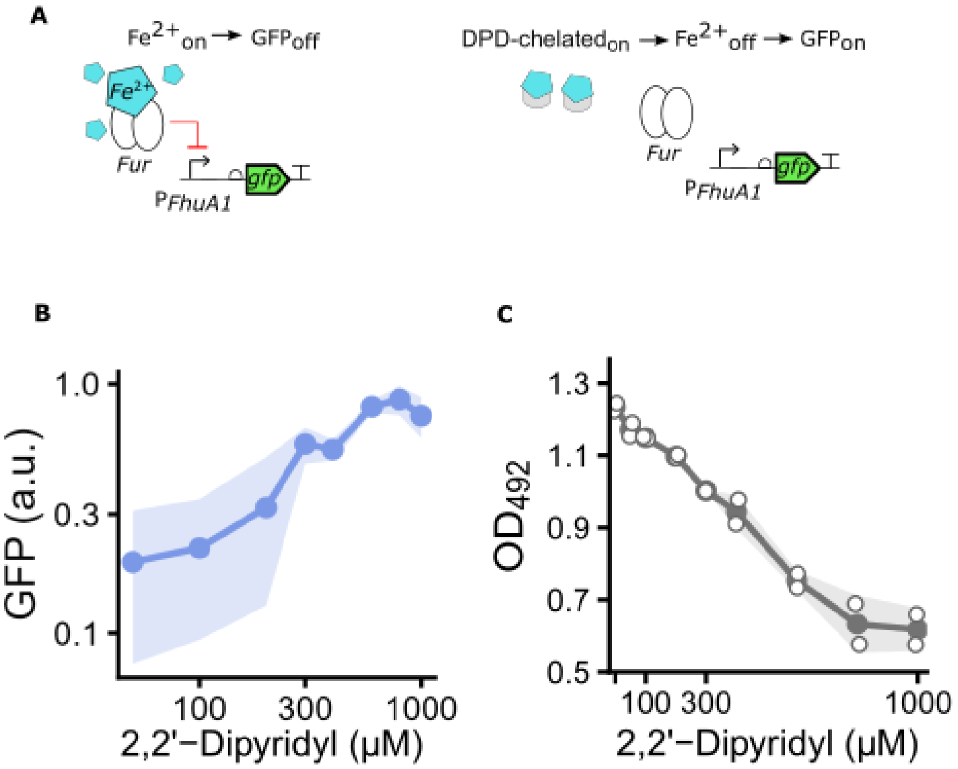


Figure 1: **Figure S1 – Effects of iron chelation by 2,2'-Dipyridyl on the $P_{FhuA1}$ -based iron sensor and growth of *P. protegens* Pf-5.** (A) Fur-mediated regulation of $P_{FhuA1}$ under iron-replete and iron-depleted conditions. In the presence of $Fe^{2+}$, Fur represses $P_{FhuA1}$, resulting in low GFP expression. Chelation of $Fe^{2+}$ by 2,2'-Dipyridyl (DPD) relieves repression and activates GFP expression. (B) GFP fluorescence after 22h̃ of incubation in LB medium supplemented with increasing DPD concentrations (0–1000 $\mu$M). (C) Endpoint $OD_{492}$ of the same cultures, showing the effect of DPD on growth. Only Pf-5 was tested. Points show independent biological replicates (n = 2) and lines represent the mean. Note that the iron titration in the main text (Figure 2B) was performed in M9-glucose, not LB.

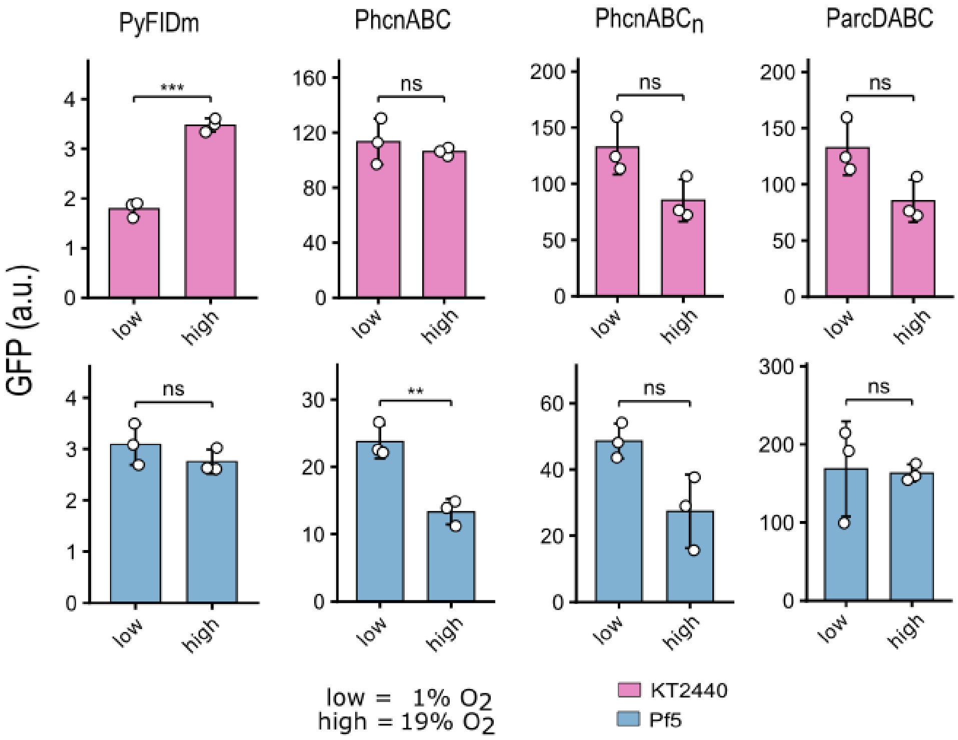


Figure 2: Figure S2 – **Screening of oxygen-responsive promoters in *P. putida* KT2440 and *P. protegens* Pf-5**. GFP output of three candidate promoters driving a gfp-reporter on the same SEVA backbone: $P_{yFIDm}$ (low-oxygen-responsive in *E. coli* ([1]), $P_{arcDABC}$ (oxygen sensing in *P. protegens* CHA0 ([2]), and $P_{hcnABC}$ (low-oxygen-responsive in *P. protegens* CHA0 ([3,4]), the latter tested with the standard RBS $P_{hcnABC}$ and with its native RBS $P_{hcnABCn}$. Cultures were measured under low (1% $O_2$) and atmospheric (19% $O_2$) oxygen. Fluorescence values were normalized to $OD_{600}$ and corrected for autofluorescence. KT2440 is shown in pink and Pf-5 in blue. Bars show the mean ± SD of three independent biological replicates (open circles). Low versus high oxygen was compared separately for each host (Welch's two-sample *t*-test): ns, not significant; **, p < 0,01; ***, p <0.001. Only $P_{hcnABC}$ in Pf-5 showed significant, correctly oriented state separation, and it was selected for the rest of the study.

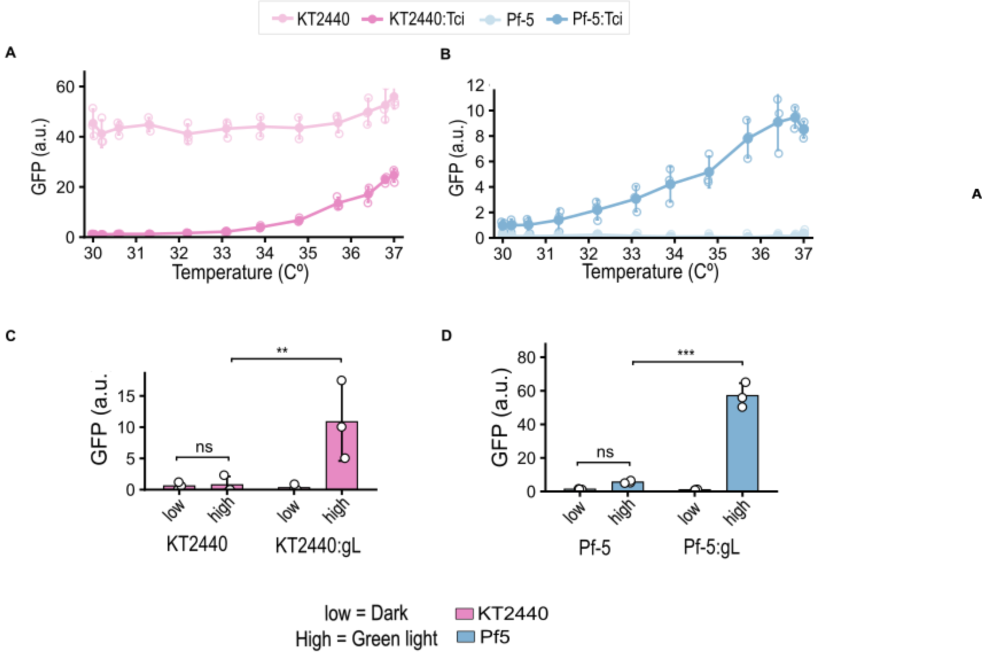


Figure 3: Figure S3. **Regulatory controls for the temperature- and green-light-responsive systems in *P. putida* KT2440 and *P. protegens* Pf-5**. Controls are parental hosts carrying the sensor plasmid but lacking the chromosomally integrated (Tn7) regulatory network module. (A–B) Temperature-dependent GFP expression in KT2440 (A) and Pf-5 (B), with and without the cI857 module, across a temperature gradient from 30 to 37 °C. Without cI857, expression is temperature-independent but at opposite levels in the two hosts: constitutively high in KT2440 and constitutively low in Pf-5. (C–D) GFP expression in the dark and under green light (520 nm) in KT2440 (C) and Pf-5 (D), with and without the CcaS/CcaR module. Green light increased fluorescence only in strains carrying the light-regulatory module (two-way ANOVA; light construct interaction: KT2440, $p = 0.0238$; Pf-5, $p < 0.001$). Fluorescence values were normalized to $OD_{600}$ and corrected for autofluorescence. Lines and bars show means, error bars indicate ±SD, and open circles represent individual biological replicates ($n = 3$).

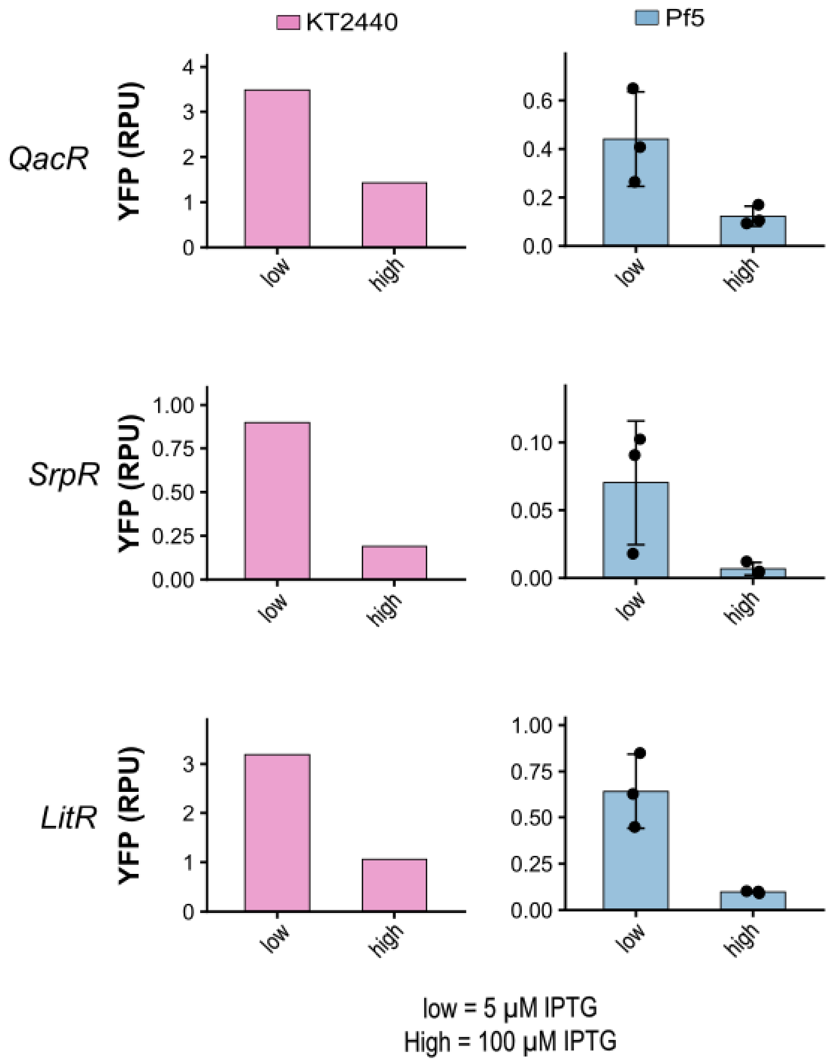


Figure 4: **Figure S4. Reference response of the QacR-, SrpR-, and LitR-based NOT gates to IPTG in *P. putida* KT2440 and *P. protegens* Pf-5.** Standalone inverter outputs under low and high IPTG (5 and 100 µM), shown as a reference for the sensor-coupled gates in the main Figure 4. The data are replotted from previous work (KT2440, Tas et al, 2021;[5] Pf-5, Rico et al. 2026[6]) and were not acquired in parallel with the circuits in Figure 4. YFP output is expressed as relative promoter units (RPU).

Table 1: List of bacterial strains used in this study

| **Strain** | **Description/Genotype** | **Reference/Source** |
|---|---|---|
| *P. putida* KT2440 | Wild-type, prototrophic derivative of *P. putida* mt-2 cured of plasmid pWW0. | [7] |
| *P. protegens* Pf-5 | Wild-type, plant-associated strain isolated from the cotton rhizosphere. | [8] |
| *P. putida* KT2440:Tci | KT2440 attTn7::[$P_{\mathrm{J23106}}$–BCD12–cI857–$T_{\mathrm{rpoCD}}$]. Temperature-sensing strain ($\mathrm{Gm}^R$). | This work; GenBank: QB285683 |
| *P. putida* KT2440:gL | KT2440 attTn7::[ccaS–ho1–pcyA–ccaR regulatory module]. Green-light-sensing strain ($\mathrm{Gm}^R$). | This work; GenBank: QB285682 |
| *P. protegens* Pf-5:Tci | Pf-5 attTn7::[$P_{\mathrm{J23106}}$–BCD12–cI857–$T_{\mathrm{rpoCD}}$]. Temperature-sensing strain ($\mathrm{Gm}^R$). | This work; GenBank: QB285685 |
| *P. protegens* Pf-5:gL | Pf-5 attTn7::[ccaS–ho1–pcyA–ccaR regulatory module]. Green-light-sensing strain ($\mathrm{Gm}^R$). | This work; GenBank: QB285684 |
| *E. coli* pir2 | $F^-$ Δlac169 rpoS(Am) robA1 creC510 hsdR514 endA recA1 uidA(ΔMluI)::pir; cloning strain expressing wild-type Pir, allowing maintenance of R6K$\gamma$-origin plasmids. | Invitrogen |
| *E. coli* TOP10 | $F^-$ mcrA Δ(mrr-hsdRMS-mcrBC) $\phi$80lacZΔM15 ΔlacX74 recA1 araD139 Δ(ara-leu)7697 galU galK rpsL($\mathrm{Str}^R$) endA1 nupG. Strain used for cloning. | Invitrogen |
| *E. coli* DH5$\alpha$ | $F^-$ $\phi$80lacZΔM15 Δ(lacZYA-argF)U169 recA1 endA1 hsdR17($\mathrm{rK}^-$ $\mathrm{mK}^+$) phoA supE44 $\lambda^-$ thi-1 gyrA96 relA1. Strain used for cloning. | Thermo Scientific |
| *E. coli* DH10B | $F^-$ mcrA Δ(mrr-hsdRMS-mcrBC) $\phi$80lacZΔM15 ΔlacX74 recA1 endA1 araD139 Δ(ara-leu)7697 galU galK $\lambda^-$ rpsL($\mathrm{Str}^R$) nupG. Strain used for cloning. | Invitrogen |
| *E. coli* HB101 (pRK600) | HB101: $F^-$ mcrB mrr hsdS20($\mathrm{rB}^-$ $\mathrm{mB}^-$) recA13 leuB6 ara-14 proA2 lacY1 galK2 xyl-5 mtl-1 rpsL20($\mathrm{Str}^R$) glnV44 $\lambda^-$; used as helper strain for triparental mating. | [9] |

Table 2: List of plasmids and genetic constructs used in this study

| Plasmid | Relevant genetic features | Backbone / resistance | Use in this work | Reference |
|---|---|---|---|---|
| **Backbone and helper plasmids** | | | | |
| pRK600 | oriV(ColE1), RK2 $mob^+$, $tra^+$ | ColE1 / RK2 helper ($Cm^R$) | Helper plasmid for triparental mating | [10] |
| pTN2 | tnsABCD transposition genes, oriV(R6K) | R6K ($Ap^R$) | Provides the transposition functions required for mini-Tn7 chromosomal integration | [11] |
| pSEVA281 | SEVA cloning vector | SEVA ($Ap^R$) | Golden Standard Level 0 backbone | SEVA collection,[12] |
| pSEVA2219-g1 | SEVA cloning vector | SEVA ($Km^R$) | Golden Standard Level 1 backbone | Golden Standard collection,[13] |
| pSEVA2219:Tn7-g1 | SEVA cloning vector containing Tn7 insertion sites | SEVA ($Gm^R$) | Golden Standard Level 1 backbone for Tn7 insertion | Laboratory collection (unpublished) |
| **Source plasmids** | | | | |
| pMA-RQ-PFhuA1 | Synthetic $P_{\mathrm{FhuA1}}$ promoter fragment | pMA-RQ ($Ap^R$) | Level 0 promoter source for iron sensing | Thermo Fisher Scientific, custom DNA synthesis |

| Plasmid | Relevant genetic features | Backbone / resistance | Use in this work | Reference |
|---|---|---|---|---|
| pMA-T-PyFIDm | Synthetic $P_{\mathrm{yfiDm}}$ promoter fragment | pMX ($Ap^R$) | Level 0 promoter source for oxygen sensing | Thermo Fisher Scientific, custom DNA synthesis |
| pMX-PhcnABC | Synthetic $P_{\mathrm{hcnABC}}$ promoter fragment | pMX ($Ap^R$) | Level 0 promoter source for oxygen sensing | Thermo Fisher Scientific, custom DNA synthesis |
| pMX-PhcnABCn | Synthetic $P_{\mathrm{hcnABC}}$ fragment containing its native RBS | pMX ($Ap^R$) | Level 0 promoter source for oxygen sensing | Thermo Fisher Scientific, custom DNA synthesis |
| pMX-ParcDABC | Synthetic $P_{\mathrm{arcDABC}}$ promoter fragment | pMX ($Ap^R$) | Level 0 promoter source for oxygen sensing | Thermo Fisher Scientific, custom DNA synthesis |
| Toggle Switch Class I | LacI/cI(ts)-based toggle-switch construct | – | Template for cI857 and $P_{\mathrm{Ls1con}}$ | Laboratory collection (unpublished) |
| pGreenL | oriRK2, ccaS, ho1–pcyA, ccaR, $P_{\mathrm{cpcG2-172}}$–msfGFP | pGSRv ($Gm^R$) | Template for components of the green-light sensing system | 8 |

| Plasmid | Relevant genetic features | Backbone / resistance | Use in this work | Reference |
|---|---|---|---|---|
| pSEVA2219:Tn7-g1:cI857 | Tn7R, $P_{\mathrm{J23106}}$–BCD12–cI857–$T_{\mathrm{rpoCD}}$, Tn7L | pSEVA2219:Tn7-g1 ($Gm^R$) | Tn7-based chromosomal integration of the temperature regulatory module | This work |
| pSEVA2219:Tn7-g1:light-module | Tn7R; $P_{\mathrm{J23106}}$–CcaS–$T_{\mathrm{rnpB}}$; $P_{\mathrm{J23108}}$–B0034–ho1–pcyA–B0015; $P_{\mathrm{J23100}}$–CcaR–$T_{\mathrm{rpoCD}}$; Tn7L | pSEVA2219:Tn7-g1 ($Gm^R$) | Tn7-based chromosomal integration of the green-light regulatory module | This work |
| **Biosensor constructs** | | | | |
| pSEVA2219: FhuA1-gfp | $P_{\mathrm{FhuA1}}$, BCD12 RBS, domesticated gfp, $T_{\mathrm{rpoCD}}$, RK2 origin of replication | pSEVA2219 ($Km^R$) | Iron-responsive fluorescent biosensor | This work (GenBank: QB285663) |
| pSEVA2219: yFIDm-gfp | $P_{\mathrm{yfiDm}}$, BCD12 RBS, domesticated gfp, $T_{\mathrm{rpoCD}}$, RK2 origin of replication | pSEVA2219 ($Km^R$) | Oxygen-responsive fluorescent biosensor | This work (GenBank: QB285667) |
| pSEVA2219: arcDABC-gfp | $P_{\mathrm{arcDABC}}$, BCD12 RBS, domesticated gfp, $T_{\mathrm{rpoCD}}$, RK2 origin of replication | pSEVA2219 ($Km^R$) | Oxygen-responsive fluorescent biosensor | This work (GenBank: QB285666) |
| pSEVA2219: hcnABCn-gfp | $P_{\mathrm{hcnABC}}$, native hcnABC RBS, domesticated gfp, $T_{\mathrm{rpoCD}}$, RK2 origin of replication | pSEVA2219 ($Km^R$) | Oxygen-responsive fluorescent biosensor | This work (GenBank: QB285664) |
| pSEVA2219: hcnABC-gfp | $P_{\mathrm{hcnABC}}$, BCD12 RBS, domesticated gfp, $T_{\mathrm{rpoCD}}$, RK2 origin of replication | pSEVA2219 ($Km^R$) | Oxygen-responsive fluorescent biosensor | This work (GenBank: QB285665) |

| Plasmid | Relevant genetic features | Backbone / resistance | Use in this work | Reference |
|---|---|---|---|---|
| pSEVA2219: ls1con-gfp | $P_{\mathrm{Ls1con}}$, BCD12 RBS, domesticated gfp, $T_{\mathrm{rpoCD}}$, RK2 origin of replication | pSEVA2219 ($\mathrm{Km}^R$) | Temperature-responsive fluorescent biosensor | This work (GenBank: QB285668) |
| pSEVA2219: cpcG2-172-gfp | $P_{\mathrm{cpcG2-172}}$, BCD12 RBS, domesticated gfp, $T_{\mathrm{rpoCD}}$, RK2 origin of replication | pSEVA2219 ($\mathrm{Km}^R$) | Green-light-responsive fluorescent biosensor | This work (GenBank: QB285669) |
| **Sensor–gate constructs** | | | | |
| pSEVA2219: FhuA1-QacR-yfp | $P_{\mathrm{FhuA1}}$–Q1 RBS–QacR inverter; $P_{\mathrm{QacR}}$–B0035–YFP | pSEVA2219 ($\mathrm{Km}^R$) | Iron-responsive NOT gate based on QacR | This work (GenBank: QB285671) |
| pSEVA2219: FhuA1-SrpR-yfp | $P_{\mathrm{FhuA1}}$–S1 RBS–SrpR inverter; $P_{\mathrm{SrpR}}$–B0035–YFP | pSEVA2219 ($\mathrm{Km}^R$) | Iron-responsive NOT gate based on SrpR | This work (GenBank: QB285670) |
| pSEVA2219: FhuA1-LitR-yfp | $P_{\mathrm{FhuA1}}$–Q1 RBS–LitR inverter; $P_{\mathrm{LitR}}$–B0035–YFP | pSEVA2219 ($\mathrm{Km}^R$) | Iron-responsive NOT gate based on LitR | This work (GenBank: QB285672) |
| pSEVA2219: hcnABC-QacR-yfp | $P_{\mathrm{hcnABC}}$–Q1 RBS–QacR inverter; $P_{\mathrm{QacR}}$–B0035–YFP | pSEVA2219 ($\mathrm{Km}^R$) | Oxygen-responsive NOT gate based on QacR | This work (GenBank: QB285675) |
| pSEVA2219: hcnABC-SrpR-yfp | $P_{\mathrm{hcnABC}}$–S1 RBS–SrpR inverter; $P_{\mathrm{SrpR}}$–B0035–YFP | pSEVA2219 ($\mathrm{Km}^R$) | Oxygen-responsive NOT gate based on SrpR | This work (GenBank: QB285674) |
| pSEVA2219: hcnABC-LitR-yfp | $P_{\mathrm{hcnABC}}$–Q1 RBS–LitR inverter; $P_{\mathrm{LitR}}$–B0035–YFP | pSEVA2219 ($\mathrm{Km}^R$) | Oxygen-responsive NOT gate based on LitR | This work (GenBank: QB285673) |
| pSEVA2219: ls1con-QacR-yfp | $P_{\mathrm{Ls1con}}$–Q1 RBS–QacR inverter; $P_{\mathrm{QacR}}$–B0035–YFP | pSEVA2219 ($\mathrm{Km}^R$) | Temperature-responsive NOT gate based on QacR | This work (GenBank: QB285676) |
| pSEVA2219: ls1con-SrpR-yfp | $P_{\mathrm{Ls1con}}$–S1 RBS–SrpR inverter; $P_{\mathrm{SrpR}}$–B0035–YFP | pSEVA2219 ($\mathrm{Km}^R$) | Temperature-responsive NOT gate based on SrpR | This work (GenBank: QB285677) |

| Plasmid | Relevant genetic features | Backbone / resistance | Use in this work | Reference |
|---|---|---|---|---|
| pSEVA2219: ls1con-LitR-yfp | $P_{\mathrm{Ls1con}}$–Q1 RBS–LitR inverter; $P_{\mathrm{LitR}}$–B0035–YFP | pSEVA2219 ($\mathrm{Km}^R$) | Temperature-responsive NOT gate based on LitR | This work (GenBank: QB285678) |
| pSEVA2219:cpcG2-172-QacR-yfp | $P_{\mathrm{cpcG2-172}}$–Q1 RBS–QacR inverter; $P_{\mathrm{QacR}}$–B0035–YFP | pSEVA2219 ($\mathrm{Km}^R$) | Green-light-responsive NOT gate based on QacR | This work (GenBank: QB285679) |
| pSEVA2219:cpcG2-172-SrpR-yfp | $P_{\mathrm{cpcG2-172}}$–S1 RBS–SrpR inverter; $P_{\mathrm{SrpR}}$–B0035–YFP | pSEVA2219 ($\mathrm{Km}^R$) | Green-light-responsive NOT gate based on SrpR | This work (GenBank: QB285680) |
| pSEVA2219:cpcG2-172-LitR-yfp | $P_{\mathrm{cpcG2-172}}$–Q1 RBS–LitR inverter; $P_{\mathrm{LitR}}$–B0035–YFP | pSEVA2219 ($\mathrm{Km}^R$) | Green-light-responsive NOT gate based on LitR | This work (GenBank: QB285681) |
| **Reference/control constructs** | | | | |
| pSEVA2219: j23101-yfp-rpu | $P_{\mathrm{J23101}}$–RiboJ10–B0064–YFP; L3S2P21 terminator | pSEVA2219 | Constitutive reference construct used for RPU normalisation | This work (GenBank: QB285686) |

Table 3: List of primers used in this study.

| Name | Sequence (5'–3') | Use | Source |
|---|---|---|---|
| PS1 | AGGGCGGCGGATTTGTCC | Verification of cloning in pSEVA plasmids | 6 |
| PS2 | GCGGCAACCGAGCGTTC | | |
| PS3 | GAACGCTCGGTTGCCGC | Verification of spurious integration of pTn7/pBAMD | |
| PS4 | CCAGCCTCGCAGAGCAGG | | |
| PS5 | CCCTGCTTCGGGGTCATT | Verification of spurious integration of pTn7/pBAMD | |
| PS6 | GGACAAATCCGCCGCCCT | | |
| GlmS-UP-FW | CTCCGCCGACTTCAGAATATC | Screening PCR to confirm Tn7-mediated chromosomal integration in *P. putida* KT2440 | |
| GlmS-DOWN-RV | GCCAATCCTGTACACCATTCC | | |
| PFL_6211_For | CGCTCTACACCTGGATCGAC | Screening PCR to confirm Tn7-mediated chromosomal integration in *P. protegens* Pf-5 | Sequence provided by Laura Cutugno (CSIC), unpublished |
| PFL_6212_Rev | ATCCTCTACACCATCCCGCT | | |
| pLs1con-GS-for | CAATGCGAGAATTCAGGTCTCTGGAGACAGATAACCATCTGCGGTG | PCR amplification of $P_{\mathrm{Ls1con}}$ promoter (Golden Standard A–B sites) | This work |
| pLs1con-GS-rev | TAGCACCAAAGCTTTGGTCTCAAGTATCCTTTGCATACCCTGCTG | | This work |
| cIts-GS-for | CAATGCGAGAATTCAGGTCTCTAATGAGGAAACCGGTTATGAGCAC | PCR amplification of cI857 regulator (Golden Standard D–G sites) | This work |
| cIts-GS-rev | TAGCACCAAAGCTTTGGTCTCAAAGCTCAGCCAAACGTCTCTTCAGG | | This work |

| Name | Sequence (5'–3') | Use | Source |
|---|---|---|---|
| AB_ccaS_for | caatgcgaGAATTCAggtctctGGAGaggaaaccggttTTTACGGCTAGCTCAGTCCT | PCR part I of the green-light module: CcaS from pGreenL (Golden Standard A–B sites) | This work |
| AB_ccaS_rev | tagcaccaAAGCTTtGGTCTCAAGTACTAAGCTCGAGGCAAATGGT | | This work |
| BC_rnpB_for | caatgcgaGAATTCAggtctctTACTaggaaaccggttTCGGTCAGTTTCACCTGATTTACG | PCR part II of the green-light module: rnpB terminator from pGreenL (Golden Standard B–C sites) | This work |
| BC_rnpB_rev | TACCCGGGaGGTCTCTATGGGACAGTCATTCATCTTTCTGC | | This work |
| CG_light-tn7-fw2 | caatgcgaGAATTCAggtctctCCATaggaaaccggttTTCAGCTTGGCTTTACCGTC | PCR part III of the green-light module: Ho1–PcyA–CcaR cassette from pGreenL (Golden Standard C–G sites) | This work |
| CG_light-tn7-rev2 | ccaAAGCTTtGGTCTCAAAGCCTAGTTTTTCCCTTGGCACAAAGATTTTTCCGTTAAATTGGCCAG | | This work |
| GSmod-QacR-For | caatgcgaGAATTCAggtctctTACTAGTCGTCAAGTGCTGTGCTTGC | PCR amplification of QacR inverter gate (Golden Standard B site) | This work |
| GSmod-SrpR-For | caatgcgaGAATTCAggtctctTACTAGCGCTCAACGGGTGTGCTTCC | PCR amplification of SrpR inverter gate (Golden Standard B site) | This work |
| GSmod-LitRL1-For | caatgcgaGAATTCAggtctctTACTAGTCATAAGTCTGGGCTAAGC | PCR amplification of LitR inverter gate (Golden Standard B site) | This work |
| GSmod-actvp-For | caatgcgaGAATTCAggtctctTACTAGCTGTCACCGGATGTGCTTTCC | PCR amplification of promoter activity vector (Golden Standard B site) | This work |
| GSmod-notgate-rev | tagcaccaAAGCTTtGGTCTCAAGCGGGACCAAAACGAAAAAAGGC | Universal reverse primer for inverter-gate amplification targeting the terminator region (Golden Standard I site) | This work |

| Name | Sequence (5'–3') | Use | Source |
|---|---|---|---|
| GS_RPU_J23101-F | caatgcgaGAATTCaGGTCTCAGGAGGATAAGTCCCTAACTTTTACAGCTAGC | PCR amplification of the RPU standard control plasmid (Golden Standard B site) | This work |
